# ProCbA: Protein Function Prediction based on Clique Analysis

**DOI:** 10.1101/2020.11.24.396432

**Authors:** A. Khanteymoori, M. B. Ghajehlo, S. Behrouzinia, M. H. Olyaee

## Abstract

Protein function prediction based on protein-protein interactions (PPI) is one of the most important challenges of the Post-Genomic era. Due to the fact that determining protein function by experimental techniques can be costly, function prediction has become an important challenge for computational biology and bioinformatics. Some researchers utilize graph- (or network-) based methods using PPI networks for un-annotated proteins. The aim of this study is to increase the accuracy of the protein function prediction using two proposed methods.

To predict protein functions, we propose a Protein Function Prediction based on Clique Analysis (ProCbA) and Protein Function Prediction on Neighborhood Counting using functional aggregation (ProNC-FA). Both ProCbA and ProNC-FA can predict the functions of unknown proteins. In addition, in ProNC-FA which is not including new algorithm; we try to address the essence of incomplete and noisy data of PPI era in order to achieving a network with complete functional aggregation. The experimental results on MIPS data and the 17 different explained datasets validate the encouraging performance and the strength of both ProCbA and ProNC-FA on function prediction. Experimental result analysis as can be seen in Section IV, the both ProCbA and ProNC-FA are generally able to outperform all the other methods.

## 1. Introduction

Proteins are large, complex, essential and the most important molecules of life. They are main constituents in all living organisms and associated the second cell weight after water to themselves. Proteins are also responsible for some of the most important functions in an organism. Defending the body from antigens, involved in muscle contraction and movement, facilitate biochemical reactions and helping to coordinate certain body activities are some of protein functionality.

In this regard, Protein Function Prediction is one of the most important fields of study in system biology and also is one of the major challenges in the Post-Genomic era. There are different methods to achieving good function prediction. These methods use distinct approaches such as sequence and structural similarity and also gene expression profiles.

Though some percent of proteins can be expected to work in relative isolation, but it is not valuable to study a protein in isolation. Protein interactions play key roles in their functionality and to perform a specific function. Interaction of proteins can have a structure such as network. This structure is called Protein-Protein Interaction Network (PPI). In this network, nodes consider as proteins and edges represent as the interactions between proteins when two proteins interact with each other. Since the structure of network can be more various, therefore the position of each protein in interaction network plays an important role for understanding the cell activities. Therefore, Protein function implications can often be inferred via PPI networks. These inferences methods are based on the fact that the function of unknown proteins find out through discovering their interaction with a known protein target having a known function. For these reasons, it is critical to develop graph-based methods to predict protein functionality [2-10].

The recent availability of protein data and importance interaction between proteins (exploiting the protein similarity) led to the development of various methods based of interaction network for predicting protein functions. Several progress reports have been published in this area take advantage of network-based methods to capture the interaction between proteins and employ them to predict protein functions.

NG et al. [11] considered the so-called protein function pair approach, is carried forward from the protein domain pair approaches. Their approach is based on Kim et al. [12, 13] by incorporating a randomization procedure in order to assign function–function correlation score for a protein function pair, which could facilitate protein function prediction [11]. In [14], Zhu et al. proposed a Semantic and Layered Protein Function Prediction (SLPFP) framework. SLPFP is an unknown protein functions predictor and a new clustering-based function prediction algorithm at different functional layers within the Function Catalogue (FunCat) Scheme and also from different clusters rather than from just one.

To addresses the issues of protein similarity measurement and prediction domain selection, Zhu et al. proposed an innovative approach to predict functions of unknown proteins iteratively from a PPI dataset. The iterative approach of [15] considers semantic similarity of protein interactions is based on the multi-layered information carried by protein functions as dynamic features of protein structure [15]. Hou et al. [16] take into account aggregating the functional correlations among relevant proteins to predict protein functions from PPI data.

This functional aggregation considers the positive impact of each relevant and negative repeated protein functions on the final prediction results [16]. Differently, Zhu et al. [17] used a functional connectivity feature to represent the strength of a protein’s impact on its neighbor’s functions. The functional connectivity approach of [17] is a PPI network-based method.

According to the above approaches, we try to propose an effective approach to predict protein functions. To this end, we focus on the protein function prediction and in this regard develop a method, called Protein Function Prediction based on Clique Analysis (ProCbA). ProCbA composes a set of interacting components that every component describes by its structure and purpose and expressed in its functioning. Major component of our proposed method is Data Preprocessing component that removes some of the additional protein-protein interactions under certain conditions. This component processes the downloaded data from MIPS before the proposed method can use them. After pre-processing stage, the proposed method applies on the data. In this method, PPI network partitions to the different size of a clique that will be used to extracting specification and average number of adjacency proteins. In the following, theses cliques are analyzed in order to functions prediction of unknown proteins.

Addition of ProCbA, we propose Protein Function Prediction on Neighborhood Counting using functional aggregation (ProNC-FA). ProNC-FA is based on Neighborhood Counting with no addition contribution. In order to achieve complete protein-protein interactions data, this method only use integration of different scientific literature and databases to functional aggregation. The evaluation results of ProCbA and ProNC-FA demonstrate the effectiveness of the proposed method and the capability of the method in providing better predictions results compared to the well-known methods.

The remainder of this paper is organized as follows. Section II describes some related researches in different aspects of our work. Protein function prediction and details of the proposed method are explained in Section III. In this section, data preprocessing is described in Part A, method of Protein Function Prediction based on Clique Analysis (ProCbA) and method of Protein Function Prediction on Neighborhood Counting using functional aggregation (ProNC-FA) are described in Part B and C. Section IV presents the obtained results. Finally, Section V concludes this paper and provides some direction for improving this method.

## 2. Materials and methods

### 2.1. Protein Function Prediction based on Clique Analysis (ProCbA)

In order to protein function prediction, this study develops ProCbA method which utilized graph theory concept. The structure and main components of ProCbA is illustrated in Figure 1. In this figure, each component denotes the key functionalities of the proposed method, and each arrow corresponds to relations between components. Preprocessing component is an important step in the ProCbA that processes its input data to produce appropriate output. Another key component in **Error! Reference source not found**. is Clique Extraction. This component identifies and extracts all the maximal cliques of the input network then sends them as input to Clique Evaluation component. In Clique Evaluation component, necessary processing and evaluation on the extracted clique in protein network is performed.

**Figure 1.**
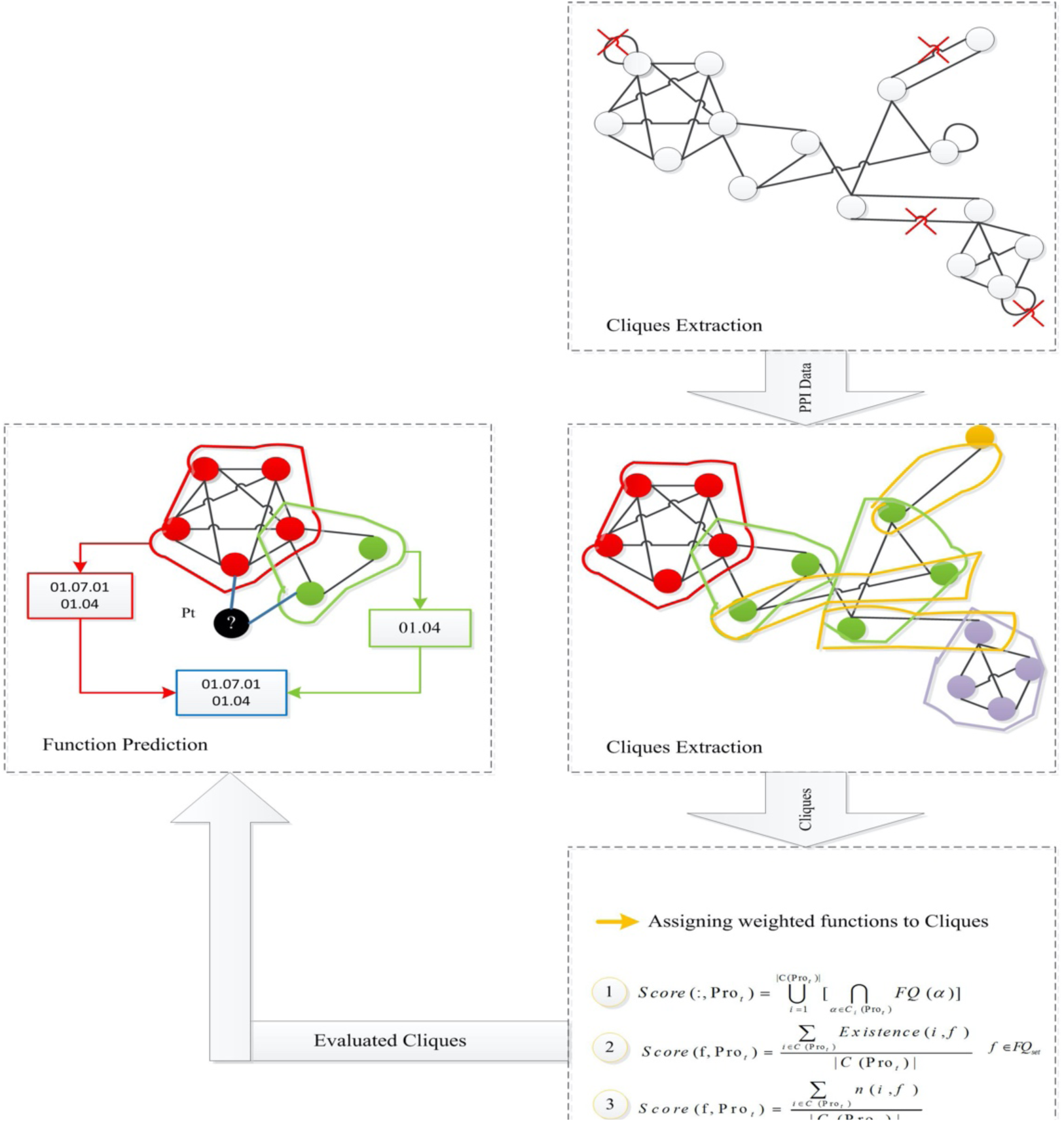
Description pf critical components of ProCbA method

The final and most important component of ProCbA is Function Prediction. This component uses two steps to choose the right protein function. In the first step of Function Prediction, number of possible functionalities of each protein identified. In the second step, this component uses two important parameters as input to protein function prediction: the first parameter is the number of functionalities of each protein and the second one is functionality frequency. In the following, each section gives more details about what component do during method execution. The ProCbA algorithm is described in Algorithm 1.

#### Algorithm 1 ProCbA: Protein Function Prediction with Clique-based Analysis

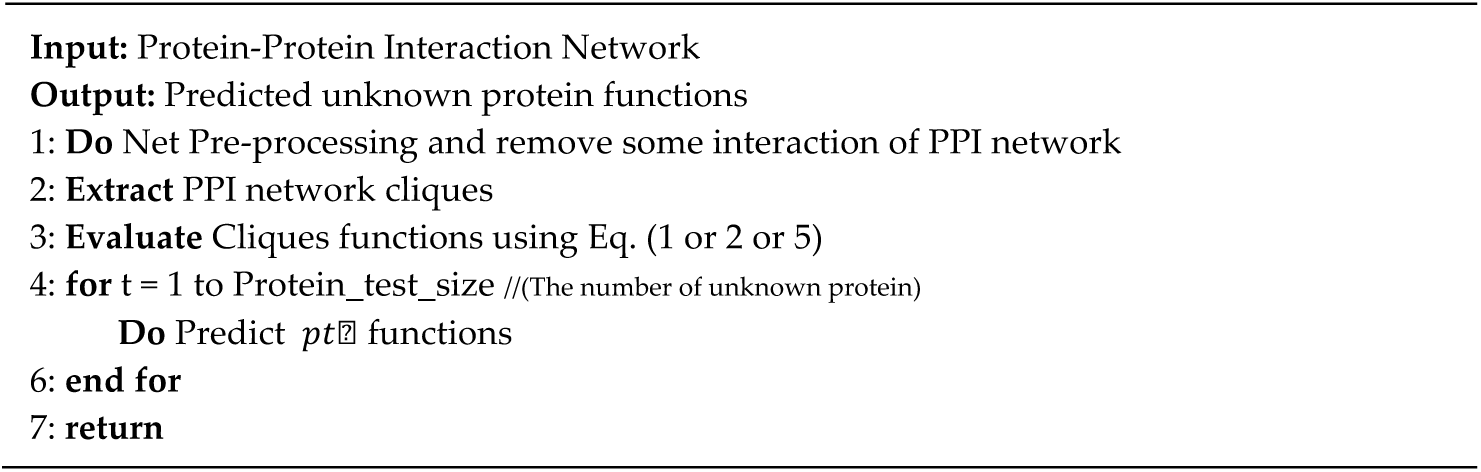

#### 2.1.1 Preprocessing

Main goal of this component is network preprocessing that removes some of the protein-protein interactions under certain conditions. This component process input data that were downloaded from MIPS before used by another component. In this component, all of the interactions that satisfy one of below conditions are removed from protein-protein interaction list. These conditions are generally as follows:

1. Remove duplicate transactions.
2. Remove the protein-protein interactions which proteins are same in interactions, or in the other words, each protein interacts herself.
3. Remove the protein-protein interactions which at least one of the proteins has not any functions.

After analysis of interaction based on 3 conditions, this component produces appropriate output. Output of this step, is a binary and symmetric adjacency matrix that is named PPI-graph or PPI-Network. PPI-graph is Protein-Protein Interaction graph based on this adjacency matrix that each elements of the matrix indicates whether pairs of vertices are adjacent or not in the graph. PPI network is an undirected graph. Figure 2 shows example of adjacency matrix.

**Figure 2.**
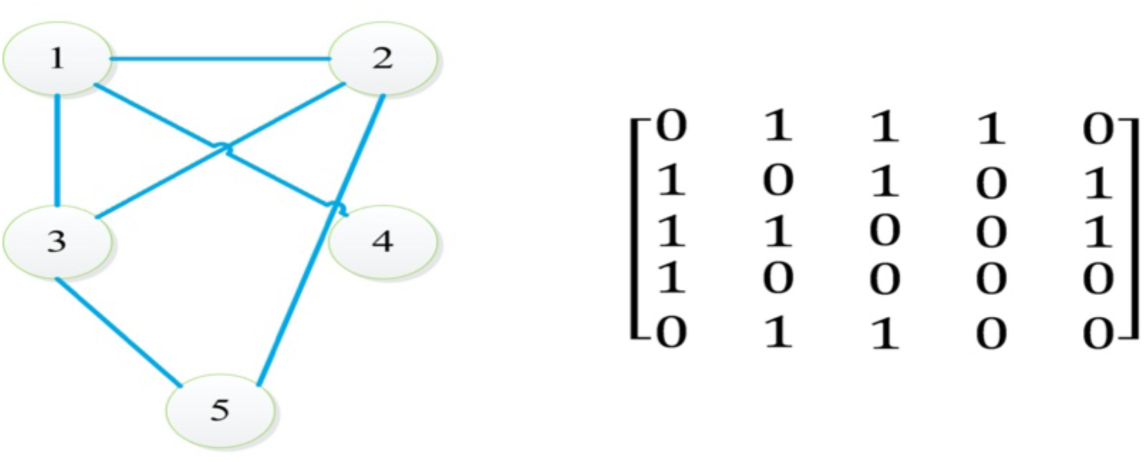
The extraction of adjacency matrix from PPI Network

**Figure 3.**
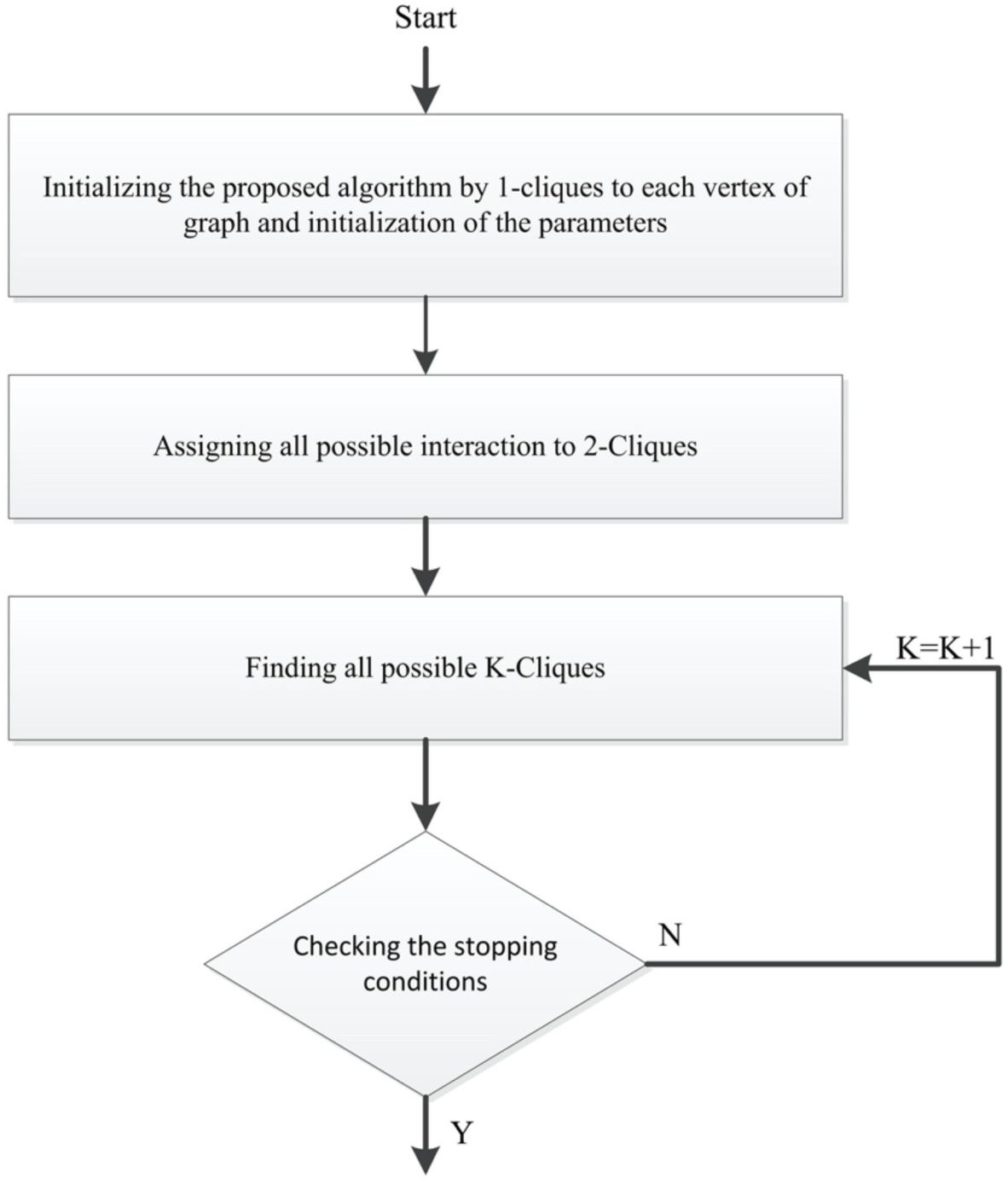
Flowchart of the Clique Extraction

**Figure 4.**
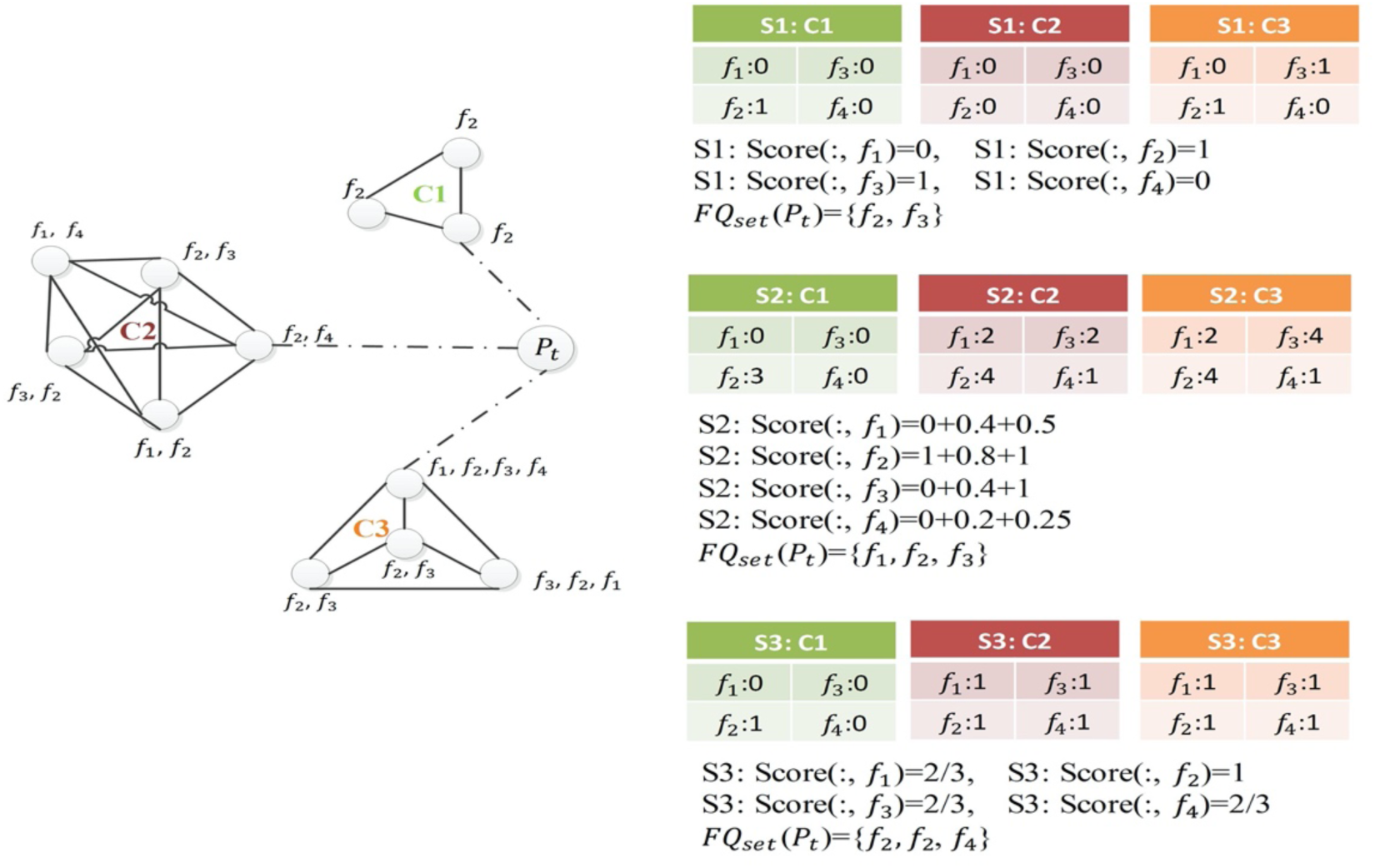
The example of demonstrating the effect of the three proposed strategies to Clique Evaluation

**Figure 5.**
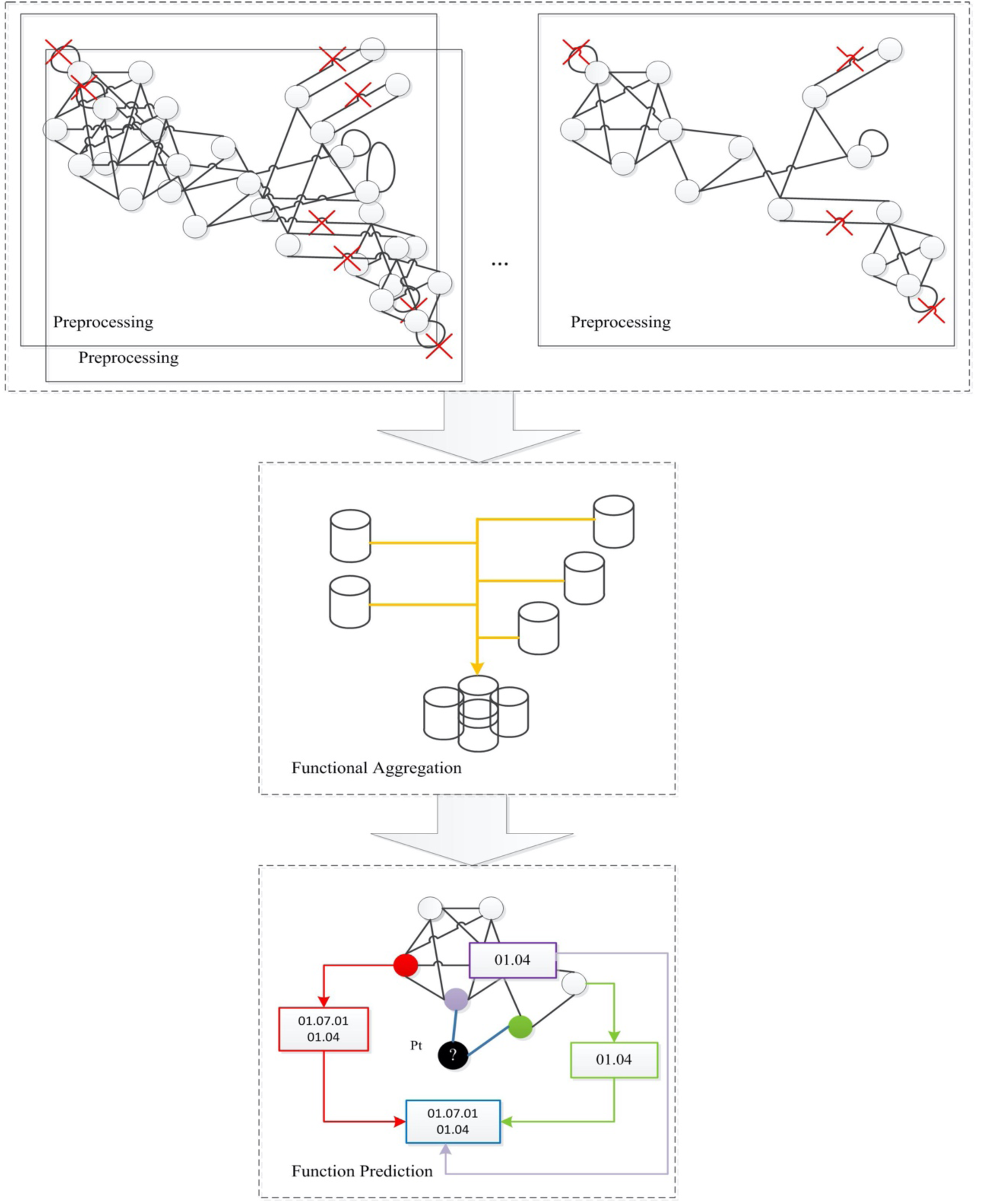
The illustration of the function prediction process of ProNC-FA

**Figure 6.**
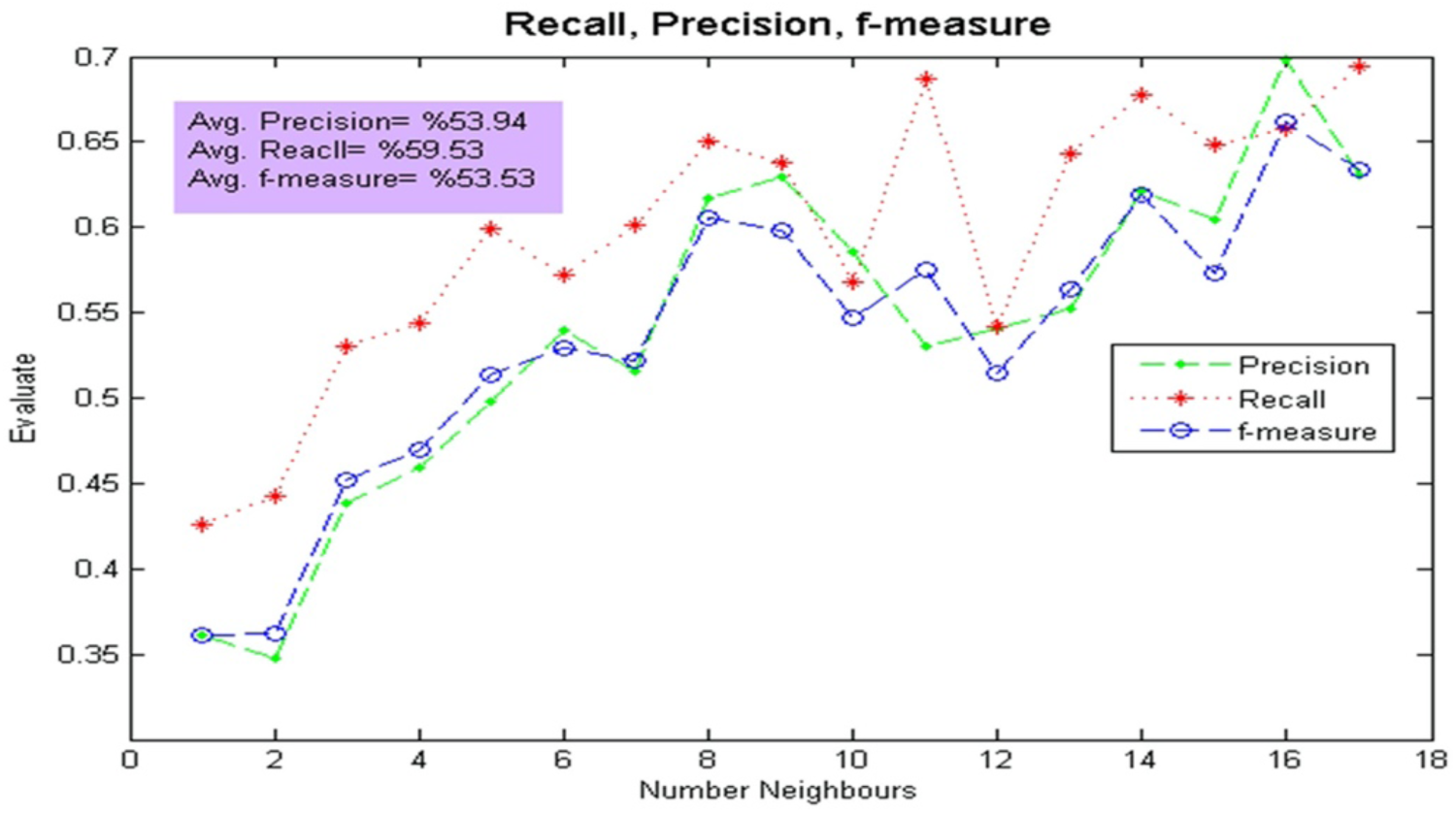
Performance of ProCbA based on neighbourhood degrees of protiens

#### 2.1.2 Clique Extraction

A clique in undirected graph *G*, is a subset of vertices that are pair-wise adjacent in the graph. Clique is one of the fundamental concepts in graph theory. Maximum clique of graph *G*, is a clique if and only if it has maximum cardinality among all possible cliques of graph G. Finding Maximum clique is an NP-hard problem [18, 19] and, for this reason, several exact algorithms have been developed for solving this problem [20].

Having the same functions between proteins can be concluded protein-protein interactions with high probability. As a result, it is concluded that, the clique is a meaningful concept from a biological perspective [21]. The used algorithm to extracting clique consists of four phases as illustrated in **Error! Reference source not found**.. After the initialization is performed, the algorithm iterates phases 3 and 4 until stopping conditions are met.

In initialization phase, since proteins can only interact with one protein, each protein is assigned to 1-clique. In phase two, all possible interaction is assigned to 2-clique. In phase 3, all proteins that can interact with one of protein in 2-cliques, are searched to finding 3-clique. This search continues until creation of all 3-cliques in graph. The phase 3 is repeated to produce all possible 4, 5, … and K-cliques. Finally, the algorithm stops if the stop conditions are met and algorithm cannot create any other maximal clique.

#### 2.1.3 Clique Evaluation

Clique Evaluation component analyzes and collects the set of adjacency cliques of each protein with unknown functions. The collected cliques set for each protein can be repeated for different cliques set but howbeit, they keep without removing repeated cliques due to their importance. Although, this approach increases the running time of the method, but as to achieve the highest prediction, accuracy is more important compared to the running time.

In the following, the goal is calculation of functions frequency. Due to this goal, all identified clique’s set are used to assign appropriate scores to all possible clique’s functions using one of the following three strategies S1, S2 and S3:

1. S1: Analysis of common functionalities to be considered

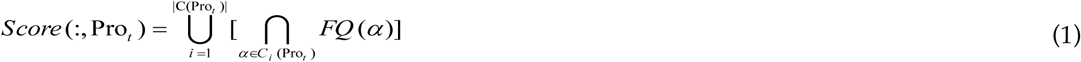

where is *C* (pro _t_) set of all adjacency cliques to test protein
2. S2: In the second strategy, one of locally scoring schemes is applied to create scored functions. This schema considers the presence probability of all functions in each clique. Continue union of all high scored functions of cliques assign to test protein. One reason of using this strategy is overcome the limitation of first’s strategy in noisy interaction. For example, when in the specific clique, all proteins have the similar functions except one protein (that has different functions), first strategy remove different functions. Removing alone functionality can be mistake strategy in some cases. Scoring function of this strategy define as:

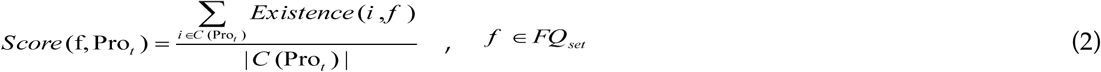 Where *pro*_*t*_ is unknown protein, *C*(*pro*_*t*_) is set of all cliques that is connected to *pro*_*t*_ where F*Q* is set of possible functions of all cliques which as follows:

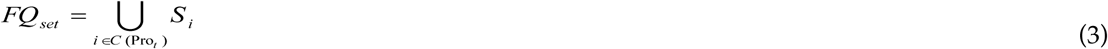

and *S* _*i*_ is union of functions of *i*^*th*^ clique as follows:

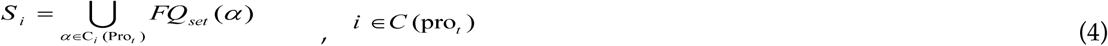
3. S3: The final strategy is somewhat similar to the second strategy. The third strategy counts the number of evidences for each possible function in each clique. Continue union of all high scored functions of cliques assign to test protein. Scoring function of this strategy define as:

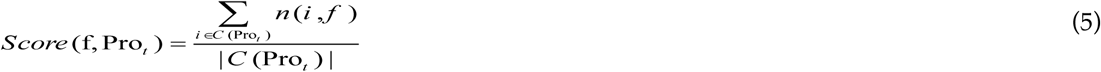

Different of three explained strategy are shown in **Error! Reference source not found**..

#### 2.1.4 Function Prediction

Assigning relative functions from candidate functions is one of the most important problems in proteomic study and protein function prediction. There is not any knowledge of the number of protein functions in the Biological Theory. As a matter of this fact, several methods have been developed to selecting number of functions and assign functionalities to an unknown protein. Selection of fixed number for all functions [22] and average numbers of functions of all network proteins are different strategies that have been used to determining number of functionalities.

To measure the number of protein functions, ProCbA uses Eq. (6) that is defined by

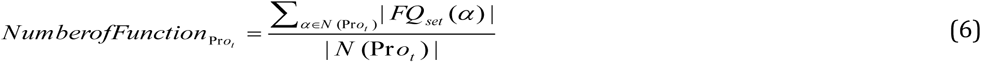

where *N* (Pr *o*_*t*_) denotes the number of test protein neighbors. Forasmuch as, Eq. (6) uses average numbers of functions of all neighbor proteins, it can be best strategy to determining number of functions.

### 2.2 Protein Function Prediction on Neighborhood Counting using functional aggregation (ProNC-FA)

Due to the tentative identification of data in proteomic study, some of the major important limitations of protein-protein interaction networks are incomplete and noisy data. To address this issue, this study introduces ProNC-FA, which takes advantage of the integrated data as the functional aggregation. This functional aggregation investigates the impact of integrated data in performance and accuracy of protein function prediction and the impact of integrated data in performance reduction of previous approach. The functional aggregation feature of ProNC-FA reduces the impact of repeated functional information on the prediction.

**Error! Reference source not found**. shows an overview of ProNC-FA method. As illustrated in **Error! Reference source not found**., ProNC-FA uses functional aggregation component based on Neighborhood Counting algorithm that is proposed by Schwikowski [22]. Goal of this component is aggregation and combination of all interaction in Protein-Protein Network. The improvement of speed and performance are the best result of ProNC-FA.

Main component of ProNC-FA is functional aggregation component that use output of preprocessing component. ProNC-FA uses 17 different PPI dataset as illustrated in **Error! Reference source not found**.. First, each of dataset is processed to produce appropriate output by preprocessing component. Second, all preprocessed data integrates as an output of functional aggregation. Final, this functional aggregation data is sent as input to Clique Extraction. Evaluation result of ProNC-FA demonstrates the effectiveness of the proposed method and the capability of the method in providing better predictions results compared with existing methods.

## 3. Result and Discussion

### 3.1 Datasets

Generally, protein function prediction methods use two important resource data, namely, protein-protein interaction and functional annotation scheme. We evaluate the ProCbA by testing their performance on the tasks of predicting protein functions on MIPS [32-33, 41-42] and FunCat3 [43].

MIPS is a database for genomes and protein sequences that is provided genome-related information by The Munich Information Center in Germany. Further information of MIPS Dataset is showed in Table 1.

**Table 1.**
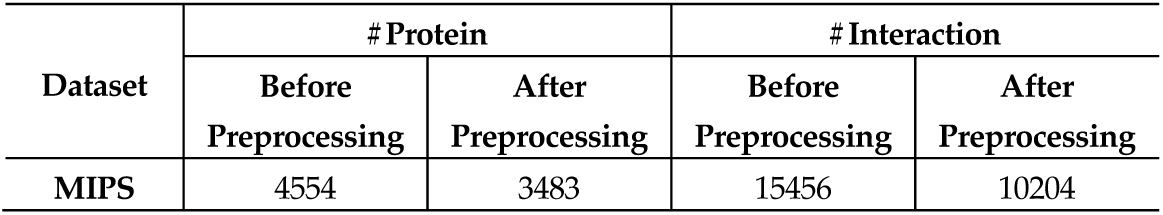
MIPS Data Specifications.

FunCat3 [30, 68] is functional annotation scheme which have wide coverage and standard hierarchical structure. The 28 existing functions in FunCat3 are organized in a hierarchical tree structure. In this study, the most informative functions of FunCat are used to testing.

In addition to what has been said, we use the 17 different datasets to achieve the complete network of Protein-Protein Interaction. The details of this used data within ProNC-FA are presented in Table 2.

**Table 2.** The specification of 17 used different dataset in ProNC-FA.

| Dataset | Before Preprocessing |  | #Protein<br>(Without interaction) | #Interaction<br>(After redundancy removal) | After Preprocessing |  |
| --- | --- | --- | --- | --- | --- | --- |
|  | #Interaction | #Protein |  |  | #Interaction | #Protein |
| Alexei [23,24] | 2238 | 1827 | 2238 | 244 | 1931 | 1519 |
| Shin [25] | 22571 | 5496 | 22568 | 1157 | 19057 | 4199 |
| Tong2004 [26] | 7941 | 2262 | 7175 | 394 | 6171 | 1812 |
| yeastHighQuality [24] | 2455 | 988 | 2455 | 32 | 2408 | 947 |
| Yeast_data_2007 [24] | 17481 | 4931 | 17194 | 974 | 14686 | 3873 |
| DIP_MMIPS_iPfam [27] | 3201 | 1681 | 2857 | 44 | 2806 | 1541 |
| Gavin2006 [28,29] | 6531 | 1430 | 6531 | 60 | 6340 | 1363 |
| Krogan [30] | 7123 | 2708 | 7084 | 317 | 6291 | 2316 |
| MINT [31] | 48321 | 5341 | 24421 | 1120 | 20527 | 4158 |
| MIPS [32, 33] | 15456 | 4554 | 12319 | 942 | 10204 | 3483 |
| Se2012 [34] | 112331 | 6012 | 112010 | 1395 | 93190 | 4612 |
| DIP2013 [35] | 22995 | 5004 | 22493 | 987 | 19697 | 3934 |
| SGD [36] | 338246 | 5999 | 222230 | 1289 | 192766 | 4710 |
| Utez [37] | 1033 | 1003 | 1033 | 165 | 700 | 764 |
| Ito2001 [38] | 4038 | 2937 | 3959 | 539 | 2984 | 2252 |
| STRING v9.1(2013) [39] | 1660496 | 6397 | 830248 | 1692 | 674911 | 4704 |
| BIOGRID2013 [40] | 338904 | 6234 | 31201 | 1523 | 192836 | 4711 |

### 3.2 Evaluation Metrics

This section presents evaluation metrics to test the effectiveness and estimate the expected accuracy of ProCbA to protein function prediction. Various evaluation metrics have been developed for evaluating effects of different prediction methods. To evaluate the proposed method, we use three evaluation metrics, namely, Precision, Recall and F-Measure.

- **Precision**: In the field of function prediction, precision is the fraction of predicted functions that are relevant to the protein. Precision criteria is defined as:

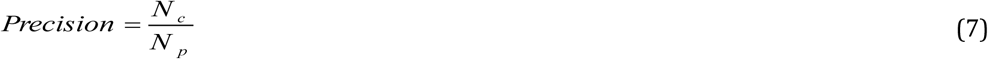 Where *N* _*p*_ is the number of predicted functions of protein and *N* _*c*_ is the number of correct predicted functions.
- **Recall**: Recall in function prediction field is the fraction of the functions that are relevant to the protein that are successfully predicted. Recall criteria is defined as:

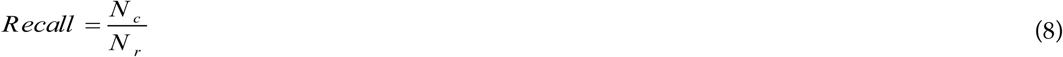 Where *N* _*r*_ is the number of identified functions of protein and *N* _*c*_ is the number of predicted functions.
- **F-Measure**: The traditional F-measure or balanced F-score is a measure that combines precision and recall is the weighted harmonic mean of precision and recall. The balanced F-Measure criteria is defined as:

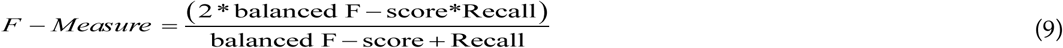 All the experiments reported in this section were performed on a system with an Intel Core i7-2410M 2.3 GHz processor with 6 GB RAM.

### 3.3 ProCbA Evaluation

Given the importance of the strategy that is used to Clique Evaluation, more specifically ProCbA prediction based on a 3 proposed strategy can be concluded different evaluation results by using the evaluation criteria expressed. Table 3 shows the results of ProCbA for various strategies. Results in Table 3 shows that the evaluation results rate depends on the used strategy. As seen in the table, the second strategy gives the best results in function prediction in three evaluation criteria and chooses to use in Clique Evaluation component in ProCbA.

**Table 3.** Dependency of prediction results on different strategies.

| Strategy | ProCbA |  |  |
| --- | --- | --- | --- |
|  | Precision | Recall | F-Measure |
| S1 | 42.93594 | 52.73269 | 43.8063 |
| S2 | 52.81869 | 57.5718 | 52.04874 |
| S3 | 25.80969 | 28.87197 | 25.85506 |

To test the ProCbA’s accuracy, we adopt k-fold cross validation. In this process, all the Protein-Protein interactions are randomly divided into k subsets. Each time of k, one subset of k subsets is selected as testing data and the rest subsets are used as the training set. In this paper, k is 10. In the following, experimental results of ProCbA on k cross validation subsets are showed in Table 4.

**Table 4.**
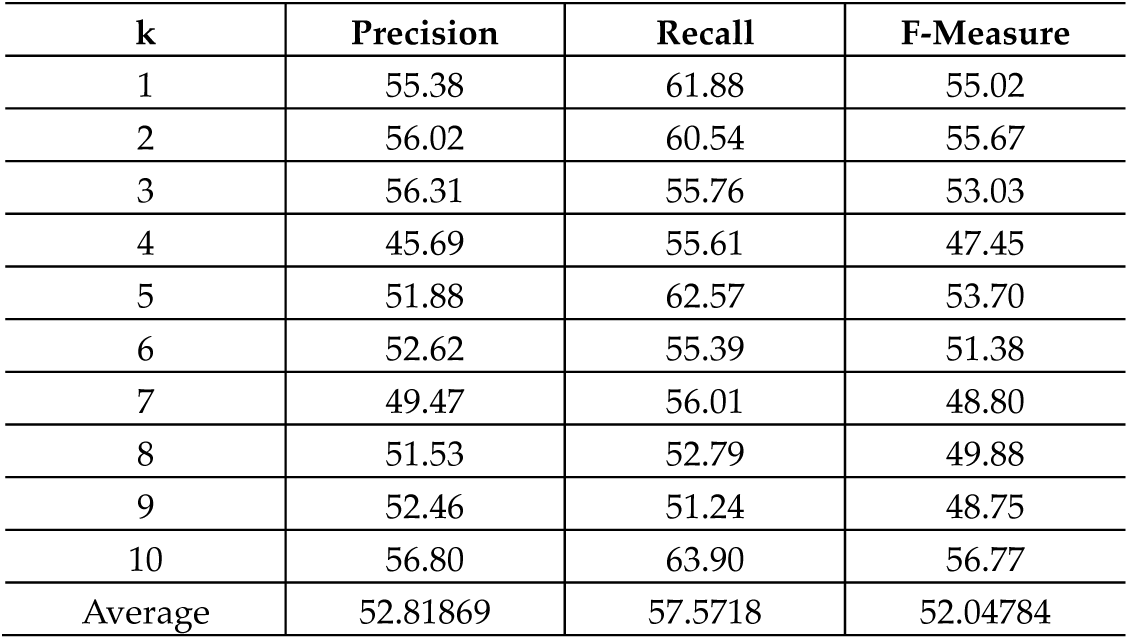
Overall precisions, recalls and F-Measure of ProCbA method on the MIPS dataset based on k-cross validation.

| k | Precision | Recall | F-Measure |
| --- | --- | --- | --- |
| 1 | 55.38 | 61.88 | 55.02 |
| 2 | 56.02 | 60.54 | 55.67 |
| 3 | 56.31 | 55.76 | 53.03 |
| 4 | 45.69 | 55.61 | 47.45 |
| 5 | 51.88 | 62.57 | 53.70 |
| 6 | 52.62 | 55.39 | 51.38 |
| 7 | 49.47 | 56.01 | 48.80 |
| 8 | 51.53 | 52.79 | 49.88 |
| 9 | 52.46 | 51.24 | 48.75 |
| 10 | 56.80 | 63.90 | 56.77 |
| Average | 52.81869 | 57.5718 | 52.04784 |

As can be seen, the mean results obtained for the different k, is 52% that is best result with different train data in each 10 steps. The reason is that, ProCbA is not sensitive to input data or input interaction network. In other words, the results show that the proposed algorithm is resistant or robust to network changing.

Table 5 gives the comparisons with the other algorithms such as GM [21], GMV, *χ*^2^, FF and FCML [46]. All reported results in Table 5 are based on the presented results by [46]. According to the obtained results, ProCbA outperforms the GM, GMV, *χ*^2^, FF and FCML on F-Measure and outperforms all of the on Precision. The most current method on function prediction focuses the improvement of one of the F-Measure or Precision criteria but ProCbA can be achieved the minimum difference on these two criteria. This may be because ProCbA uses Eq. (6) to measure the functions number of proteins. The Eq. (6) can present the best estimation of the number of allocable functions to proteins.

**Table 5.** The comparison of ProCbA with five well-known algorithms.

| Algorithms | The Mean of Precision | The Mean of F-Measure |
| --- | --- | --- |
| GM | 30.69 | 29.04 |
| GMV | 31.13 | 22.41 |
| $\chi^2$ | 14.80 | 07.60 |
| FF | 28.01 | 27.05 |
| FCML | 54.83 | 43.74 |
| ProCbA | 52.81 | 52.04 |

By comparing the results of the assessment, according to what is shown in the table, ProCbA have better result in the both Precision and F-Measure. For example, the reason is that the GM algorithm only considers directed neighbor but ProCbA does not limit itself to considering directed neighbor. The neighborhood or the protein degree concludes the number of protein neighbors. In the other words, the neighborhood degree of a protein p in a protein network N is the induced the number of subnets of N consisting of all protein adjacent to p. The most important goal of this study is the proof of relationship between the neighborhood degrees of proteins with the number of proteins functions. **Error! Reference source not found**. is shown the performance of ProCbA based on neighborhood degrees of proteins.

According to increasing the number of protein neighbors, the growth of ProCbA performance in Precision, Recall and F-Measure will be more significant. In Protein-Protein Networks, with increasing and decreasing the neighborhood degree, the number of proteins is gradually decrease and increase respectively. Due to this specification of network, accuracy and performance of ProCbA are not good.

Incomplete functional annotation scheme is another consequence that can be concluded from this evaluation. Due to **Error! Reference source not found**., it is expected that the performance of algorithm increases with increasing the number of neighborhood degree. But, as can be seen in the results of the evaluation, the performance of ProCbA has oscillated from the neighborhood degree of 2. In the other words, Accuracy of ProCbA not only not improved but also has been reduced.

Interaction of proteins with proteins that have similar specific functions is an important general hypothesis in protein-protein interaction network. In this regard, two probabilities can be considered. First, general hypothesis is not correct and second, data related to functional annotation scheme is not complete. Second probability is more powerful than first. It is expected, if protein’s interacting partners identify, not only the accuracy of the proposed algorithm also the accuracy of most existing algorithms will increase. To proof of this theory, second proposed method named as ProNC-FA is presented. According to this theory, ProNC-FA uses the functional aggregation to protein function prediction.

### 3.4 ProNC-FA Evaluation

Since using the functional aggregation data is the main novelty of ProNC-FA, in this section the performance of this method using one of the first presented methods in this scope is evaluated. Table 6 demonstrates the results of ProNC-FA with 17 different datasets. Each row of this table shows the result of PPI Network made using the intersection from first row’s data till this row’s data. The column 2 and 3 in table represent the number of interactions and the number of proteins, respectively and the column 4 and 5 in table represent the min and max neighborhood degree of protein, respectively. For example, there are proteins that have interaction with one protein or 3188 proteins in the first row. Difference between the neighborhood degree of column 4 and 5 conclude two main results. First, the network is still incomplete and second, there are many pseudo interactions in networks which the difference causes an increase in neighborhood degree of some of proteins.

**Table 6.**
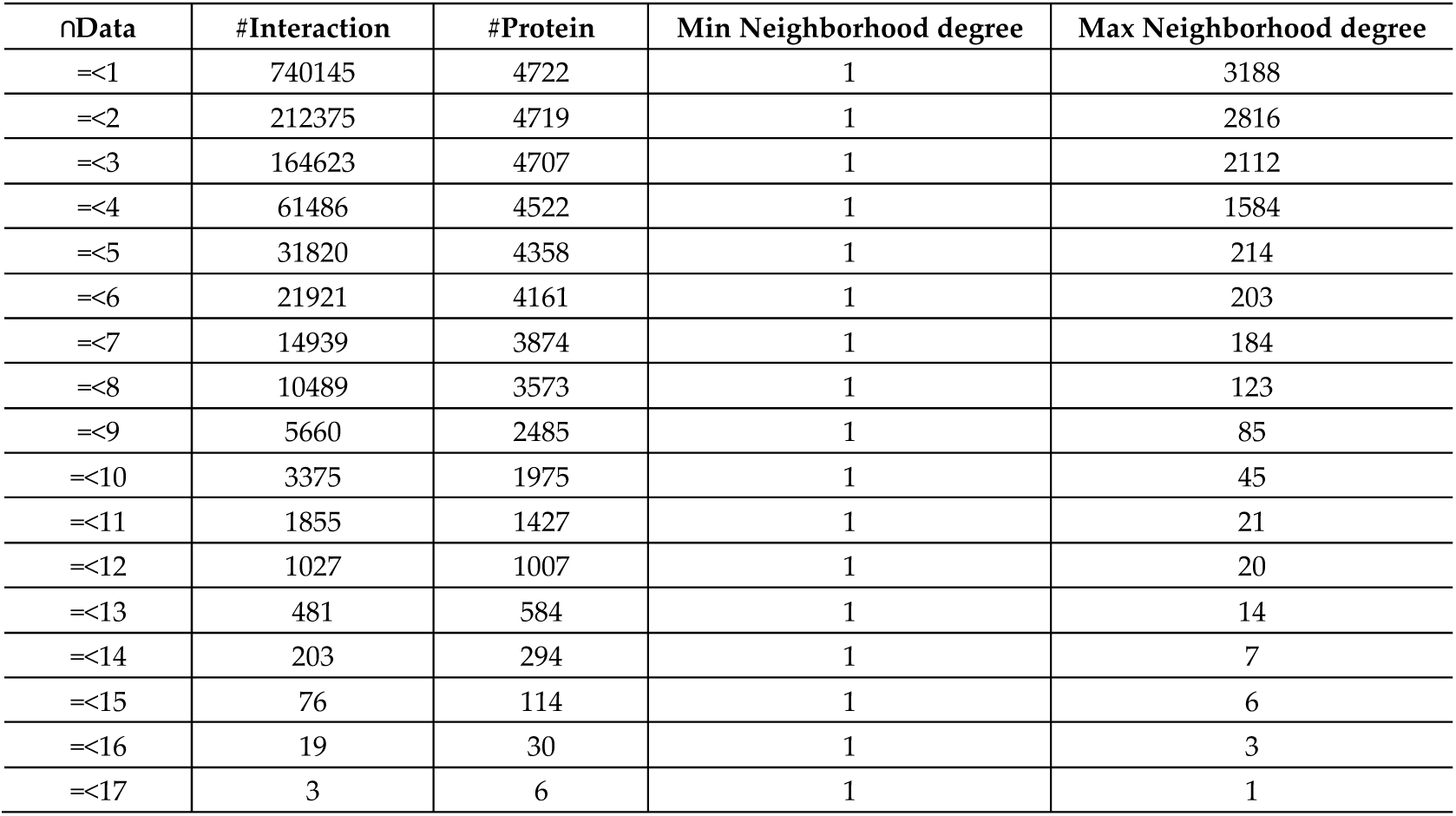
The integration results of ProNC-FA on the 17 different datasets that is used to functional aggregation.

| nData | #Interaction | #Protein | Min Neighborhood degree | Max Neighborhood degree |
| --- | --- | --- | --- | --- |
| =<1 | 740145 | 4722 | 1 | 3188 |
| =<2 | 212375 | 4719 | 1 | 2816 |
| =<3 | 164623 | 4707 | 1 | 2112 |
| =<4 | 61486 | 4522 | 1 | 1584 |
| =<5 | 31820 | 4358 | 1 | 214 |
| =<6 | 21921 | 4161 | 1 | 203 |
| =<7 | 14939 | 3874 | 1 | 184 |
| =<8 | 10489 | 3573 | 1 | 123 |
| =<9 | 5660 | 2485 | 1 | 85 |
| =<10 | 3375 | 1975 | 1 | 45 |
| =<11 | 1855 | 1427 | 1 | 21 |
| =<12 | 1027 | 1007 | 1 | 20 |
| =<13 | 481 | 584 | 1 | 14 |
| =<14 | 203 | 294 | 1 | 7 |
| =<15 | 76 | 114 | 1 | 6 |
| =<16 | 19 | 30 | 1 | 3 |
| =<17 | 3 | 6 | 1 | 1 |

According to the obtained results, small percentage of the interactions about 0.0004% is in the difference dataset and this suggests that interactions have different degree of reliability. Considering different degree of reliability as weighted protein interaction network can lead to future research.

Furthermore, the proposed method is compared with some popular methods such as PClustering[44] and PRODISTIN[45]. These methods were evaluated by Ashish on 2013[44]. Table 7 shows the obtained results from the comparison of PClustering, PRODISTIN and ProNC-FA methods on the 27 explained proteins in [44]. In general, as can be seen in the results, the ProNC-FA shows an appropriate and high accuracy compared with other methods. For example, ProNC-FA can predict all functionality of protein 19 and 27. In the most rows of Table 7, the prediction accuracy of ProNC-FA is equal to or greater than other algorithms. Using of simplest approach without any math complexity to predicting protein function is one of most important features of ProNC-FA.

**Table 7.**
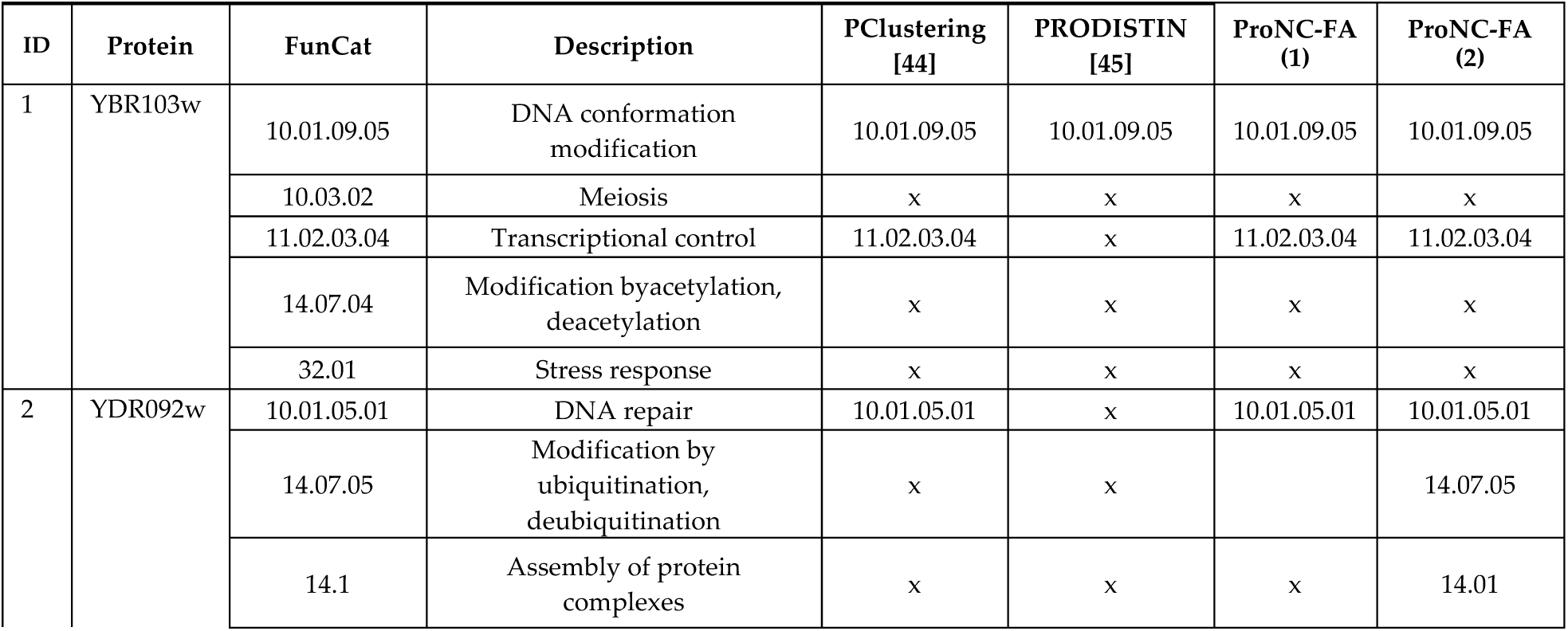
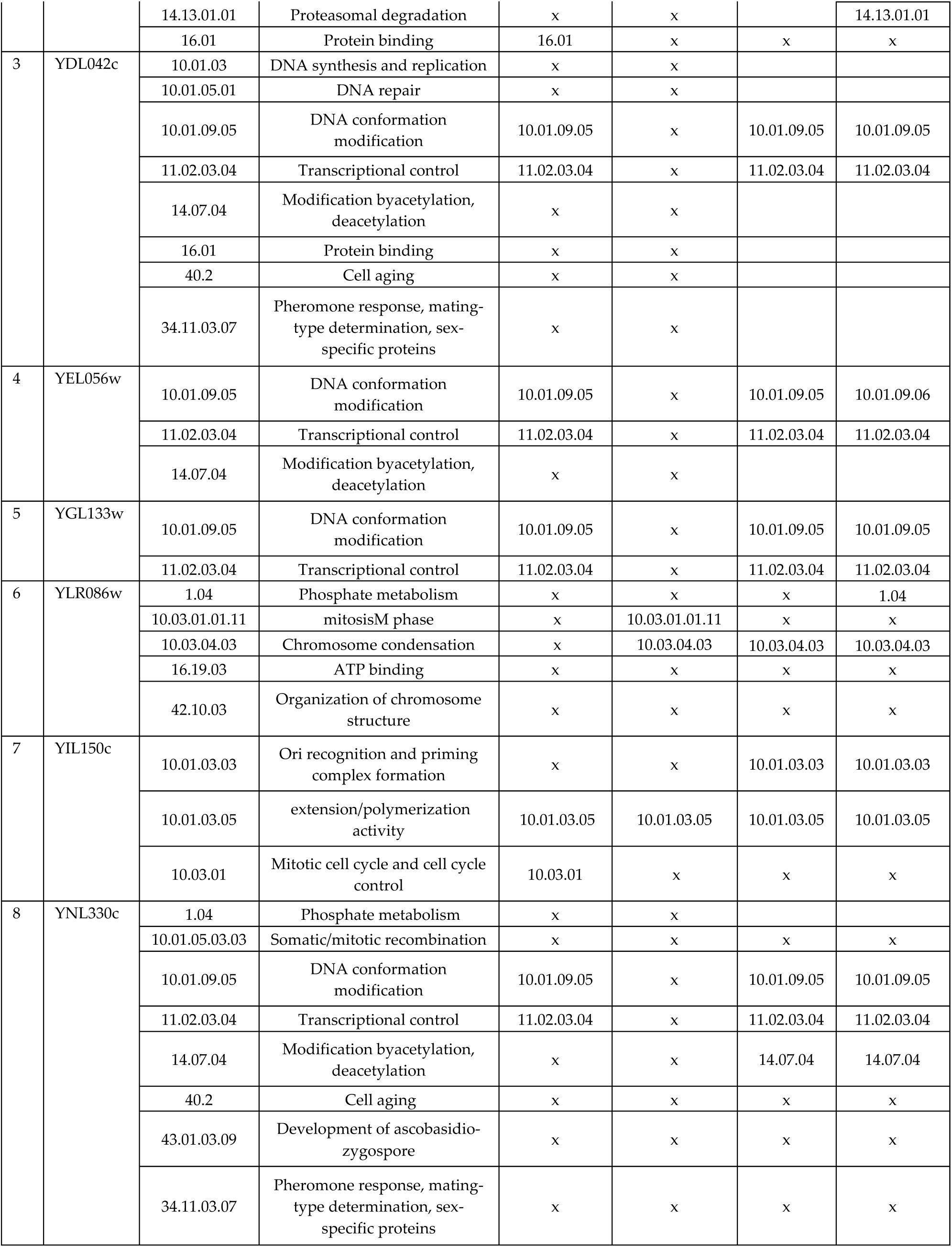
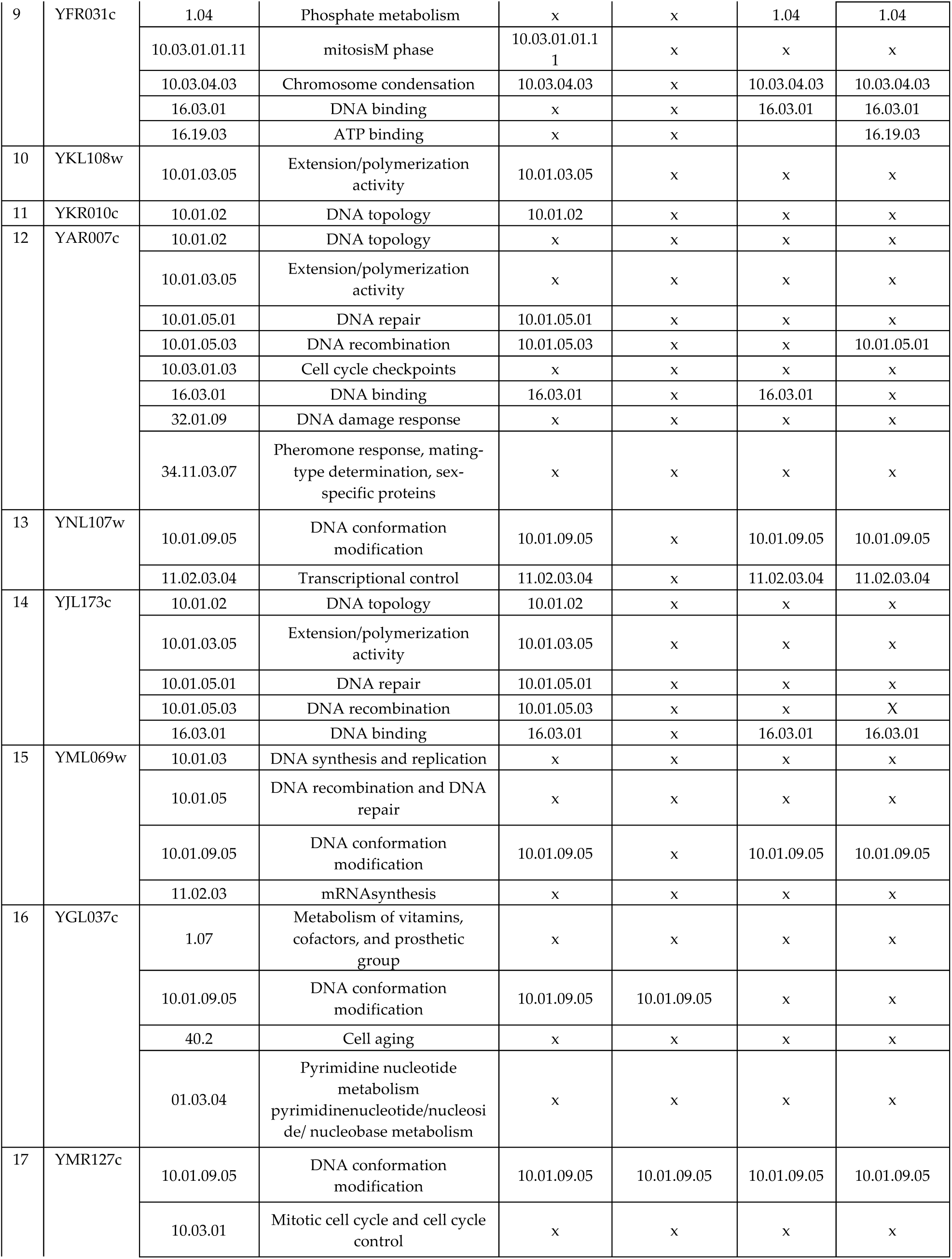
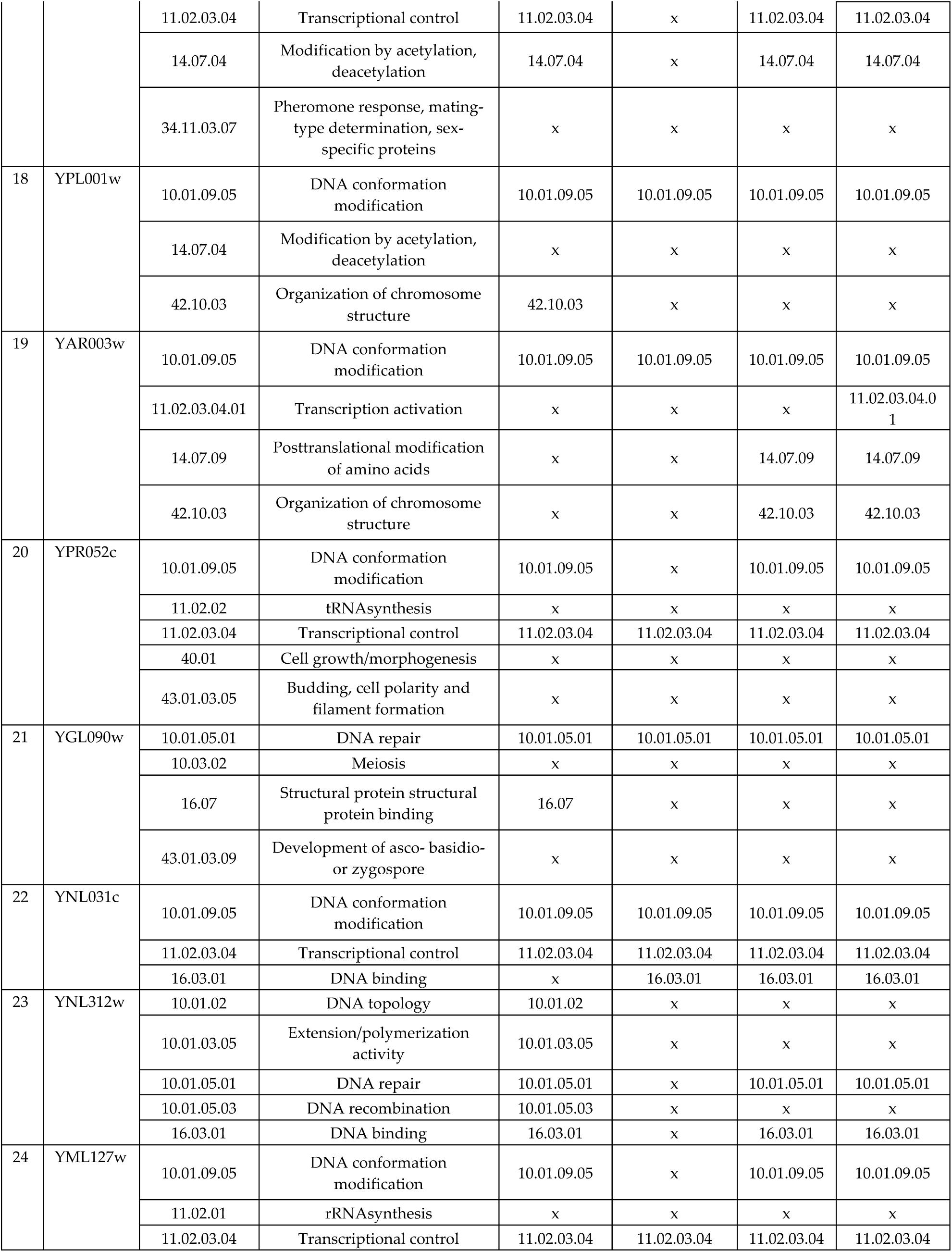
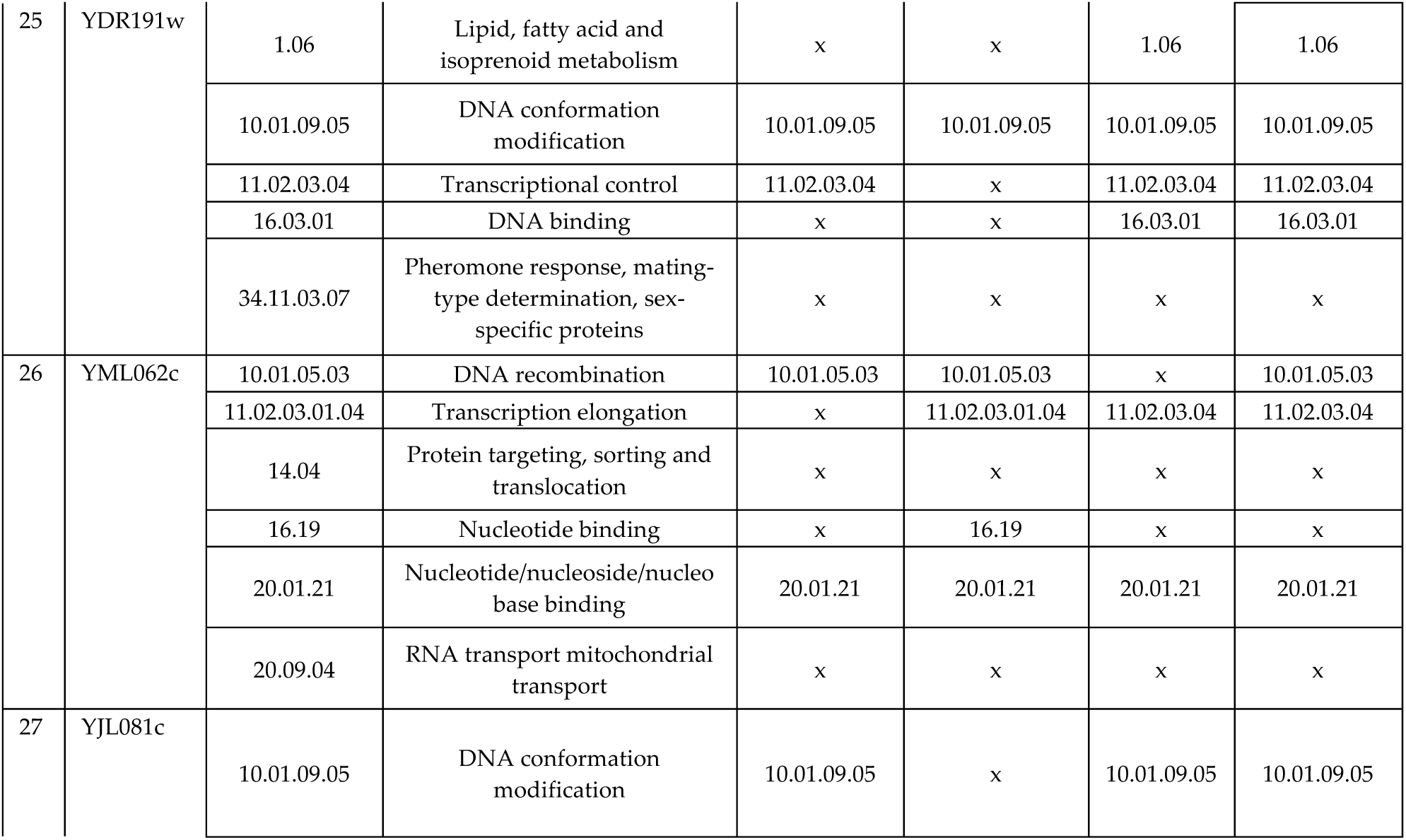
Evaluation results of ProNC-FA compared to PClustering and PRODISTIN. (ProNC-FA(1) is related to original Neighbourhood Counting algorithm with functional aggregation, ProNC-FA(2) is related to using Neighbourhood Counting algorithm based on Eq. (6) to measure the function number of proteins.

Since using the neighborhood counting method is the function prediction component of ProNC-FA, in this section, the low performance of neighborhood counting using two important proteins is evaluated and compared with ProNC-FA. Table 8 demonstrates the results of the neighborhood counting algorithm and ProNC-FA with 2 different proteins. According to the obtained results, ProNC-FA has more complete data than the neighborhood counting algorithm and can predict YBL072c protein functions with high accuracy. Although, initial information for YAL012w protein in FunCat is complete, ProNC-FA does not have acceptable result in function prediction.

**Table 8.** The benefit and drawbacks of using ProNC-FA(1) to function prediction of two known protein namely, YAL012w and YBL072c.

| Protein | # Neighbors | FunCat | Prediction Based on ProNC-FA(1) | %Frequency |
| --- | --- | --- | --- | --- |
| YAL012w | 572 | 01.01.06.05.01.01<br>01.01.09.03.01 | 01.04 | 8 |
|  |  |  | 01.07.01 | 7 |
|  |  |  | 01.05 | 7 |
| YBL072c | 606 | 12.01.01 | 12.01.01 | 26 |
|  |  |  | 11.04.01 | 12 |
|  |  |  | 16.03.03 | 11 |

Figure 7 gives the comparisons results of different algorithms such as ProCbA, ProNC-FA(1) and ProNC-FA(2). According to the obtained results, both ProNC_FA(1) and ProNC–FA(2) outperform ProCbA with high accuracy in protein interaction prediction. It should be noted that ProNC-FA has complete information of neighbors of proteins.

**Figure 7.**
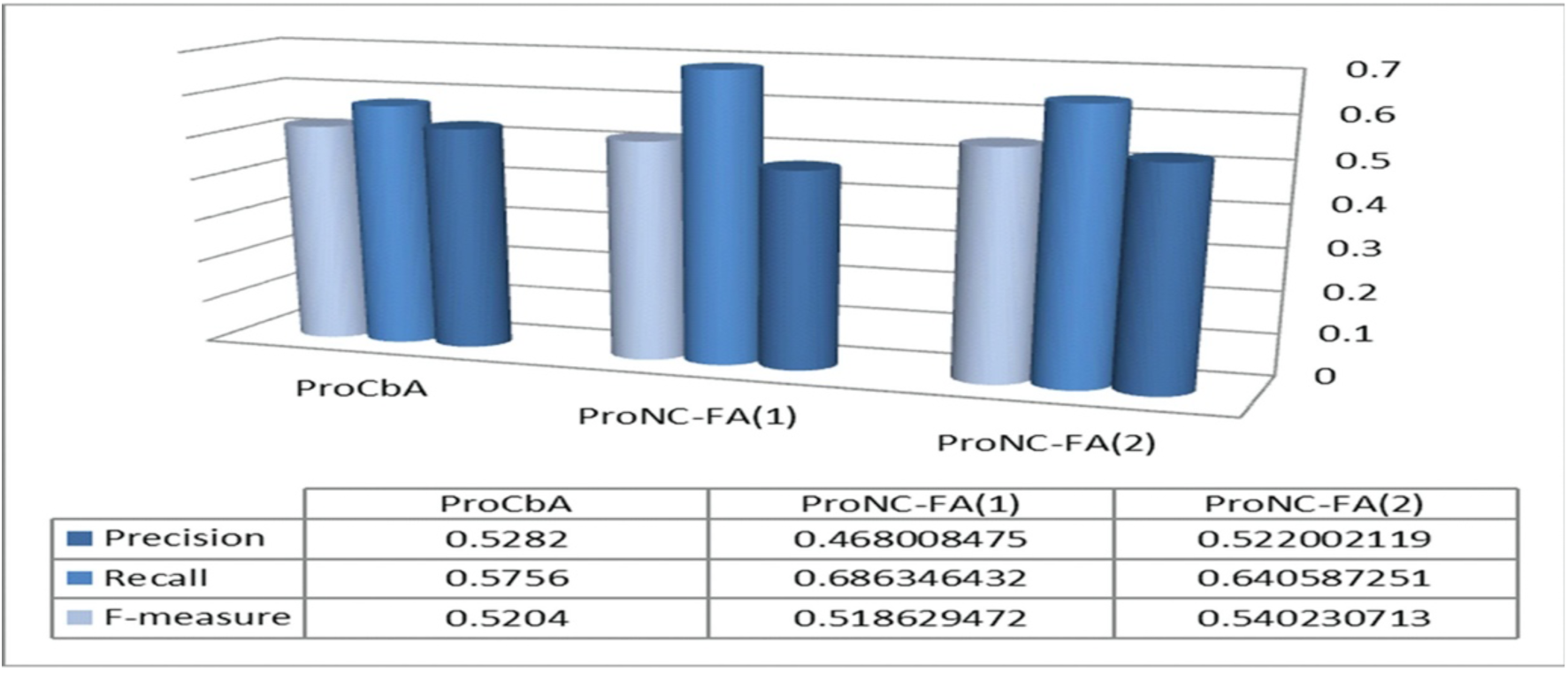
The evaluation results of proposed methods

## 4. Conclusion

Though numerous studies and many interaction detection methods have presented on protein function prediction, extracting useful knowledge can still be important and one of the challenging tasks on function prediction in the bioinformatics field. Clique based analysis is one of the most important methods. ProCbA which is presented in this study is Clique based Analysis method for protein function prediction. The evaluation results of ProCbA suggest that the cliques found by ProCbA algorithm are consistent with the biological knowledge.

One of the important problems in Protein data set is the presence of noise in data. Thus, affects reduction of noise in a data set may conclude the best result in the function prediction process. In this paper, a method based on functional aggregation data using Neighbourhood Counting algorithm named as ProNC-FA is proposed to solve this problem. In order to analysing ProNC-FA, 17 datasets as a popular benchmark is used to functional aggregation. Comparison with other algorithms is shown that the proposed method prepares promising results for function prediction.

